# Should a few null findings falsify prefrontal theories of conscious perception?

**DOI:** 10.1101/122267

**Authors:** Brian Odegaard, Robert T. Knight, Hakwan Lau

## Abstract

Is activity in prefrontal cortex (PFC) critical for conscious perception? Major theories of consciousness make distinct predictions about the role of PFC, providing an opportunity to arbitrate between these views empirically. Here we address three common misconceptions: i) PFC lesions do not affect subjective perception; ii) PFC activity does not reflect specific perceptual content; iii) PFC involvement in studies of perceptual awareness is solely driven by the need to make reports required by the experimental tasks, rather than subjective experience *per* se. These claims are incompatible with empirical findings, unless one focuses only on studies using methods with limited sensitivity. The literature highlights PFC’s essential role in enabling the subjective experience in perception, *contra* the objective capacity to perform visual tasks; conflating the two can also be a source of confusion.

## An opportunity to empirically resolve some enduring controversies

Our theoretical understanding of the neural mechanisms of conscious perception remains primitive. To make progress, investigators have begun emphasizing specific, testable hypotheses. One promising topic that has seen renewed interest is the role of prefrontal cortex (PFC) in consciousness. This empirical question can be directly addressed by experiments, providing an opportunity to arbitrate between several theoretical frameworks (Dehaene and Naccache, 2001; Lamme, 2003, 2006, 2010; Dehaene et al., 2006, 2011; de Gardelle and Kouider, 2009; He and Raichle, 2009, 2010; Kouider et al., 2010; Cleeremans, 2011, 2014; Dehaene and Changeux, 2011; Dehaene, 2014; Koch et al., 2016a, 2016b; Sandberg et al., 2016; Tononi et al., 2016). For example, while the Local Recurrency theory predicts that PFC activity is not critical for perceptual consciousness (Lamme, 2006), the Higher Order (Lau and Rosenthal, 2011) and Global Workspace (Baars, 1997, 2005; Dehaene and Naccache, 2001; Dehaene, 2014) theories assert that PFC activity plays an important causal role in enabling conscious perception. Thus, this is currently one of the more tractable and meaningful issues in the field.

## The clinical neuropsychology of consciousness: ‘classical’ case studies

Based mainly on old lesion studies (e.g., Hebb and Penfield, 1940; Fulton, 1949; Mettler, 1949; Brickner, 1952), often mentioned indirectly via secondary source (e.g., Pollen, 1999), the role of prefrontal cortex in consciousness has long been debated (Tse et al., 2005; Kouider et al., 2007; Lau and Rosenthal, 2011; Noy et al., 2015). Several investigators have called this topic into question again recently (Tsuchiya et al., 2015; Koch et al., 2016a, 2016b; Sandberg et al., 2016), asserting that PFC plays a negligible role in consciousness.

However, important details about the extent of lesions in older work are often overlooked, leading to false conclusions. For example, Koch et al. (2016a) cite work on one patient with an “almost complete bilateral frontal lobectomy” (p.319, referencing Brickner, 1952) to provide evidence that lesions to PFC do not impair consciousness. While Brickner (1952) claims that most of Patient A.’s frontal lobe is missing, simple visual inspection of the patient’s brain image reveals that massive portions of right PFC remain intact (Figure 1).

**Fig. 1.**
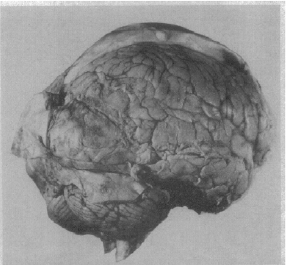
The brain of Patient A. following ‘bilateral’ frontal lobectomy (from Brickner, 1952). Patient A. was reported as the first human subject to undergo ‘bilateral’ frontal lobectomy. While this patient has previously been described as a classic example of how lesions to PFC do not impair consciousness (Koch et al., 2016a), simple visual inspection of this patient’s post-mortem brain image reveals that massive amounts of residual right PFC remained following the surgery. Therefore, the fact that this patient exhibited signs of consciousness following surgery is not surprising. Note also that this patient also had a large posterior meningioma exerting pressure on extrastriate cortices.

Indeed, much older work on psychosurgical patients often cited in discussions of PFC (Hebb and Penfield, 1940; Fulton, 1949; Mettler, 1949) centers on cases with only unilateral or partial removal of frontal cortex, or polar frontal and orbitofrontal resection. For example, Hebb and Penfield (1940) stress a remarkable lack of debilitating symptoms in their patient that had extensive bilateral removal of frontal cortices. While they emphasize that their postoperative report may underestimate the amount of tissue extracted during their procedure, their diagram of the cytoarchitectonic fields mapped by Brodmann (their Fig. 8) shows that much of Broca’s area, superior frontal gyrus, and posterior middle gyrus remained following surgery, which can account for the preserved visual and limited cognitive abilities in the patient. Furthermore, Hebb and Penfield acknowledge only removing approximately one third of the frontal lobes in their summary (p. 438, 1940). Similarly, the patients described by Mettler (pp. 64-77, 1949) are also marked by partial removal of frontal cortical areas. Thus, mischaracterizations of historical data can lead to incorrect conclusions about the role of PFC in consciousness.

Importantly, there are in fact cases where large, near-complete bilateral lesions to the dorsal and ventral lateral PFC fully abolish normal conscious interaction with the external world. One of the authors (RTK) has personally tested patients with such lesions, and often they appear to be almost completely disconnected from - and unaware of - the external world. However, these cases are rarely reported in detail in published scientific articles because such patients are often unable to participate in formal neuropsychological testing (Mettler, 1949). But these patients, similar to patients with bilateral anterior cingulate lesions, often represent the classic clinical observations of severe abulia and perseveration (Barceló and Knight, 2002; Stuss and Knight, 2013), indicating a loss of capacity for conscious interaction with the environment.

Critically, regarding this general capacity to interact with and be aware of the external world, lesions to areas posterior to the frontal lobes tend to cause minimal, if any, impairments. For example, bilateral occipital lesions (Fig. 2A) can cause cortical blindness, and bilateral parietal lesions (Fig. 2B) can result in impaired visual feature binding (Friedman-Hill et al., 1995) among other visual impairments associated with Balint’s syndrome. However, while these patients have prominent sensory deficits (e.g., in the visual domain), they retain the capacity for being alert, aware and interactive with their surroundings. Similarly, extensive bilateral orbitofrontal lesions cause extensive social regulation deficits (Beer et al., 2003, 2006; Perry et al., 2016), but leave patients cognizant of sensations and fully interactive, albeit often excessively. However, bilateral lateral PFC lesions (Fig. 2D) frequently leave patients abulic, akinetic, unable to consciously process stimuli from various sensory modalities, and unable to have goal-directed interactions with the environment.

**Fig. 2.**
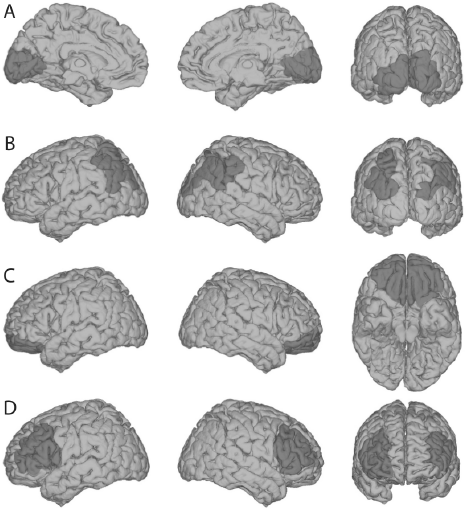
Relationships between cortical lesions and general awareness of the external world. Shown here are schematic depictions of typical lesion locations for each of the following types of impairments. (A) Bilateral occipital lesions can cause cortical blindness (Melnick et al., 2016), but preserve an individual’s general awareness of the surrounding world, via other sensory modalities. (B) Bilateral parietal lesions can cause “Balint’s Syndrome,” typically characterized by an inability to perceive multiple objects in the visual field coherently, but again, this leaves basic sensory awareness intact. (C) Bilateral orbitofrontal lesions cause problems with social regulation, but not sensory perception or consciousness. (D) Patients with bilateral lateral PFC lesions are ‘conscious’ in the sense that they are ‘awake’ - as opposed to being in a coma - but they lack the ability to engage meaningfully with the external world with a normal level of conscious awareness. Note that extensive unilateral PFC lesions do not tend to cause these symptoms, highlighting that one functional side of PFC may be sufficient to support general conscious behavior. However, further psychophysical evidence indicates that even unilateral PFC lesioned patients may suffer from specific impoverishment of subjective perceptual experiences (see below; Del Cul et al., 2009; Fleming et al., 2014).

## How do lesions to PFC specifically affect subjective perception?

The above discussion highlights that claims about consciousness based on individual case studies can be problematic due to incomplete documentation and limited data, often difficult to access. Fortunately, more recent investigations with carefully controlled psychophysical methods (Del Cul et al., 2009; Fleming et al., 2014) have investigated how, in representative groups of patients, even incomplete PFC lesions can specifically impact subjective perceptual experiences. Similar to short-term impairments induced by transcranial magnetic stimulation (TMS) (Rounis et al., 2010), incomplete lesions to PFC do not fully abolish objective visual task performance capacities, which may explain some previous null findings. But also like TMS, they can render subjective ratings of perception less meaningful (i.e., less diagnostic of actual task performance) (Rounis et al., 2010), by up to 50% (Fleming et al., 2014) (Figure 3). This observed effect was not due to an impairment of general ability for introspection, as it was specific to perception but not memory (Fleming et al., 2014); the specificity of this result was predicted independently based on another study comparing the individual variability of structural correlates for memory and perceptual metacognition in dozens of healthy subjects without brain lesions (McCurdy et al., 2013). Likewise, in another study it has been found that the effect of incomplete PFC lesions was more pronounced for subjective ratings than for objective task performance (Del Cul et al., 2009).

**Fig. 3.**
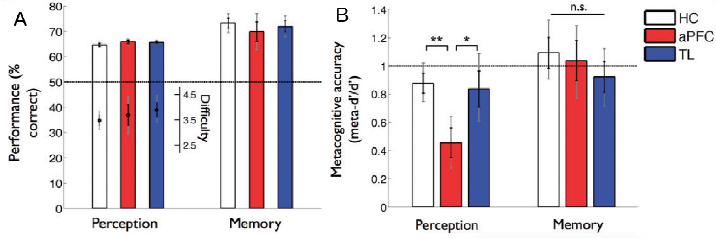
Lesion to PFC specifically impairs subjective perceptual introspection. In a study by Fleming et al. (2014), healthy controls (HC), patients with lesions to anterior PFC (aPFC), and patients with temporal lobe lesions (TL) were tested in a perception task and a memory task. (A) Overall task performance was equivalent across the three different groups in both tasks. (B) Displayed here (meta-d’ / d’) is a measure of the meaningfulness of subjective confidence ratings; i.e., the degree to which they distinguish between correct and incorrect judgments, expressed on a scale where ‘1’ implies statistical optimality and ‘0’ indicates chance-level trial-by-trial correspondence between confidence and accuracy (Maniscalco and Lau, 2012). This measure showed a nearly 50% decrease in patients with aPFC lesions, compared to patients with temporal lesions and healthy controls, and this effect was specific to perception rather than memory.

Perhaps these studies (Del Cul et al., 2009; Fleming et al., 2014) are written off as idiosyncratic examples, given that both unilateral lesions and disruption by stimulation to lateral PFC tend not to consistently impair objective visual task performance, and this means they should not affect subjective experience, either. For example, in one recent review (Koch et al., 2016a), the authors argue that subjective and objective measures of conscious perception should conform to each other, when “report protocols are implemented *judiciously*” (p.308, italics added here). However, the distinction between subjective and objective measures is central to the scientific study of consciousness (Merikle et al., 2001; Dienes, 2008; Giles et al., 2016); there is a large literature on how these measures of conscious perception systematically dissociate, and why we need to carefully consider each of them independently (for a review, see Lau, 2008; Lau and Rosenthal, 2011).

Importantly, although objective measures of task performance are often favored because they are easy to quantify, they have relatively limited face validity when it comes to studies of conscious perception. It is well known that blindsight patients (Weiskrantz, 1997) exhibit above-chance objective task performance even though they lack subjective awareness. In this context, that PFC lesions lead to selective impairment of subjective but not objective measures of perception highlights the specific and important roles played by PFC in consciousness. It would be peculiar to write this off, given the current purpose.

Perhaps some may find even a 50% impairment (Fleming et al., 2014) unimpressive, compared to, for example, the effect of lesions to V1, which can often completely abolish conscious visual phenomenology (Weiskrantz, 1997; Melnick et al., 2016). However, it is important to note that PFC functions differently from sensory cortices. For instance, neuronal coding in PFC is relatively distributed and shows a high degree of mixed selectivity (Mante et al., 2013; Rigotti et al., 2013). Therefore, to produce a large disruption of perceptual experience, many neurons distributed throughout massive regions of PFC may need to be involved. However, when PFC lesions are large and bilateral, because these same regions support many central cognitive functions (Duncan and Owen, 2000; Miller, 2000; Badre and D’Esposito, 2009; Niendam et al., 2012; Passingham and Wise, 2012), the patients may be so generally impaired that testing them immediately after extensive bilateral PFC damage is often difficult (Mettler, 1949; Knight and Grabowecky, 1995).

Also, the frontal and parietal cortices are densely connected (Barbas and Mesulam, 1981; Petrides and Pandya, 1984; Andersen et al., 1985; Cavada and Goldman-Rakic, 1989; Miller and Cohen, 2001; Croxson et al., 2005). After localized unilateral lesions to PFC, it is known that other parts of the frontal-parietal network can dynamically reorganize to compensate functionally (Voytek et al., 2010). As such, large areas spreading through both frontal and parietal areas may need to be damaged to produce consistently strong effects (Nakamura and Mishkin, 1986). This is compatible with the fact that even for classic ‘textbook’ PFC functions such as working memory, reported studies involving unilateral PFC lesions sometimes only find subtle or null effects (Mackey et al., 2016). The magnitude of the effects of PFC lesions on subjective awareness in perception (Del Cul et al., 2009; Fleming et al., 2014) should be considered within the broader context of how lesions to this region impair other cognitive functions in general.

Congruent with the above argument is the fact that chemical inactivations of PFC can sometimes lead to stronger effects, presumably because in these studies the subjects are typically tested right after the PFC is inactivated, leaving little room for alternative circuitries to take over to compensate. For example, one recent study has found that in monkeys, chemical inactivation to the prefrontal cortex impairs memory metacognition (Miyamoto et al., 2017). While this may seem on the surface to be in conflict with Fleming et al.’s (2014) result regarding a lack of memory metacognitive deficits after PFC lesions, one interpretation is that immediately after PFC lesion human patients may show impairment in memory metacognition too, but over time this function recovered due to activity in the intact remaining side of PFC, or the precuneus, which also supports memory metacognition (McCurdy et al., 2013; Richter et al., 2016). In any case, nothing takes away the fact that at the time of testing (Fleming et al., 2014), the metacognitive impairment was specific to perception, not memory, in these patients.

In rodents, it has also been shown that chemical inactivation of orbital PFC selectively impairs subjective confidence judgments without changing olfactory discrimination performance (Lak et al., 2014). In monkeys, chemical inactivation of the pulvinar, specifically the central dorsal region which likely projects to the PFC (Romanski et al., 1997; Shipp, 2003; Pessoa and Adolphs, 2010), also impairs subjective confidence judgments in a visual task, without changing task performance (Komura et al., 2013).

Overall, these results concerning subjective confidence and introspection are congruent with the classical neuropsychological findings discussed in the above section. If a perceptual object is registered in the subject’s mind with no meaningful sense of certainty whatsoever, it is unsurprising that the subject would fail to act consciously and voluntarily in reaction to the relevant object. Conceptually, it is also unclear whether conscious experiences regarding a perceptual object can ever occur without any introspectable sense of certainty that something is presented to us (Lau and Rosenthal, 2011). This is not at all to say that subjective confidence is equivalent to conscious perception, only that the latter entails at least some degree of the former.

Finally, there are also numerous studies showing that TMS and lesion to human PFC can affect many other aspects of visual perception (Barceló et al., 2000; Turatto et al., 2004; Ruff et al., 2006; Philiastides et al., 2011; Lee and D’ Esposito, 2012; Ritzinger et al., 2012; Chiang et al., 2014; Rahnev et al., 2016), and more recent neuropsychological work has demonstrated that individuals with unilateral damage to PFC often display pronounced deficits on visual tasks. For example, Barceló et al. (2000) showed that individuals with unilateral focal lesions in the lateral prefrontal cortex exhibited impairments in the ability to regulate dependent activity in extrastriate regions and detect visual targets in the visual field contralateral to the damage. Analysis of event-related potentials (ERP) revealed that prefrontal damage reduced stimulus-evoked P1 activity in parietal and temporal-occipital areas, and abolished N2 and P3 activity generated in temporal-parietal cortices, providing critical evidence for prefrontal modulation of visual processing. Indeed, other studies have also demonstrated reduced ERP amplitudes in visual cortices following damage to PFC (Voytek et al., 2010). Thus, suggestions that PFC manipulations do not cause disturbance to conscious perception are grossly incompatible with empirical facts.

## Does PFC activity reflect specific perceptual content?

The subjective measures of conscious perception we emphasized above concern subjective visibility and confidence, which are important aspects of conscious perception; these are specifically impaired in blindsight (Weiskrantz, 1997; Persaud et al., 2011; Ko and Lau, 2012; Giles et al., 2016). But to be visually conscious, one also needs to be conscious of specific visual content. It has been argued that because PFC activity does not represent such specific content (Koch et al., 2016a), its role in consciousness is probably limited.

Theoretically, this may not be a critical issue here because in enabling conscious perception, PFC may ‘work together’ with early visual areas where the specific content resides (Lau and Rosenthal, 2011). But importantly, the claim that PFC activity does not reflect specific perceptual content is also empirically false.

The impression that PFC activity may not reflect specific perceptual content is likely due to a selective reading of the fMRI literature, where many studies adopt traditional univariate ‘activation’ approaches. But univariate fMRI is known to have limited sensitivity. Indeed, it has been noted previously that visual content may only be decoded effectively from PFC when careful analysis and advanced modeling strategies are applied (Stokes, 2015; Ester et al., 2015).

Using multivariate ‘decoding’ analyses, Wang et al. (2013) has reported that PFC fMRI pattern activity reflects specific perceptual content, under conditions where the perceptual information was ambiguous, unstable, or needed to be maintained over time. Using similar methods, Cortese et al. (2016) has recently reported that perceptual content can be decoded above chance-level from PFC even in a simple perceptual decision task. In a somewhat less direct demonstration, Hebart et al. (2016) also reported that the ‘decision variable’ supporting specific perceptual decisions can be decoded from PFC fMRI pattern activity. Additionally, recent work has shown that when subjects viewed natural movies, at least some PFC voxels can contribute to decoding the content of the percept, such as whether or not a person was in a scene in a given moment during the movie (Figure 8, Huth et al., 2016).

Perhaps one may be tempted to dismiss these studies by pointing out that the results were often statistically weak. For example, Cortese et al.’s above-chance decoding of perceptual decision from PFC did not survive Bonferroni correction (2016, Figure 3C). Such may be the nature of the limited sensitivity of fMRI decoding in these brain regions, given that coding at the neuronal level in PFC may be complex (Mante et al., 2013; Rigotti et al., 2013). Another recent meta-analysis showed that PFC fMRI decoding accuracy is limited in general, even for tasks where known physiology suggests that relevant neural representations exists in the PFC (Bhandari et al., 2017).

But we also have to be aware that vision researchers often do not pay as much attention to PFC as opposed to visual areas. Whereas retinotopic mapping methods are often carefully applied to delineate early visual areas, for PFC, researchers often use statistically inefficient ‘searchlight’ methods based on brain images imperfectly aligned across participants. Sometimes, PFC data are not even acquired or analyzed. Other times positive results are reported without much emphasis. For example, Albers et al. (2013) reported positive PFC decoding findings for visual short-term memory and imagery, but only in the supplementary results section. All of these factors may contribute to the impression of the relative paucity of positive evidence regarding whether fMRI activity in PFC reflects specific perceptual content.

Fortunately, we are not limited to fMRI studies. There are numerous single- and multi-unit recording studies in non-human primates, clearly demonstrating that specific perceptual decisions are represented in PFC (Kim and Shadlen, 1999; Mante et al., 2013; Rigotti et al., 2013). Overall, these studies are compatible with the view that PFC plays a key role in ‘reading out’ encoded perceptual information from sensory cortices, which is an important step in the perceptual process (Heekeren et al., 2004; Philiastides et al., 2011; Szczepanski and Knight, 2014).

One may argue that such decisions usually concern only two or a few categories, and do not reflect fine-grained perceptual details. But the important point here is not that they necessarily determine all the details of the perceptual content. Rather, these decisions are central parts of the perceptual process itself (Green and Swets, 1966; Ratcliff, 1978); they are *not* ‘post-perceptual’ cognitive decisions. In terms of neurophysiology, it is unclear how early sensory activity on its own can lead to conscious percepts; decisional mechanisms are needed to determine whether such activity is driven by external input, or is just spontaneous noise. Accordingly, it has long been argued that these mechanisms contribute to the subjective percept itself (de Lafuente and Romo, 2006), and they have also been linked to specific perceptual illusions (Jazayeri and Movshon, 2007). So even if PFC activity does not on its own fully determine perceptual content, it certainly reflects *some* aspects of the content of conscious experience.

## Is PFC Activity Only Related to Explicit Perceptual Reports?

The last point may not be immediately convincing to some critics of PFC theories of consciousness, as the studies mentioned all involved the subjects’ explicit reporting of perceptual content. It has been claimed that in these studies, activity in PFC does not reflect conscious perception *per se;* rather, it is confounded by the task demand to report the stimulus (Tsuchiya et al., 2015; Koch et al., 2016a). Such claims are inspired by null findings in recent studies not requiring subjects to explicitly make these reports (Frassle et al., 2014; Pitts et al., 2014; Brascamp et al., 2015; Tsuchiya et al., 2015).

Proponents of PFC theories are generally aware of this possible criticism, because this line of inquiry is not new (Lumer and Rees, 1999; Tse et al., 2005; Kouider et al., 2007). However, in at least one previous study involving no explicit reports (Lumer and Rees, 1999), positive results were indeed found in PFC. In another recent study on dreams (Siclari et al., 2017), gamma band EEG activity localized in the PFC has been linked to subsequent recall of dream experiences, even though while sleeping, the subjects did not make immediate reports to experimenters. The authors argue that such gamma activity was less reliable compared to low frequency activity not localized in PFC, but it is unclear if this is simply due to the general and well-known signal-to-noise ratio difference between high versus low frequency scalp EEG signals (Juergens et al., 1999; Niedermeyer and da Silva, 2005; Buzsaki, 2006). Nevertheless, despite these positive findings, the question remains as to why some studies failed to find positive results when demands for report were removed.

Again, the abovementioned issues of limited sensitivity and relatively imprecise analysis methods typically used in fMRI studies, specifically for univariate analysis concerning the PFC, are relevant here. Using alternative methods such as direct intracranial electrophysiological recording in human surgical epileptics (electrocorticography, or ECoG), activity related to perceptual awareness in PFC has been reported even when subjects were not required to respond to the stimulus (Noy et al., 2015). Also, the above-mentioned study regarding multivoxel decoding of movie content also showed that at least some PFC activity is involved (Figure 8, Huth et al., 2016), even though subjects viewed the movies passively, without making explicit reports.

More importantly, in direct neuronal recordings in monkeys, it has also been shown that PFC activity reflects specific perceptual content, even when the animal was viewing the stimulus passively (Panagiotaropoulos et al., 2012). One may argue that even under passive viewing, an over-trained animal may still be preparing a report even though it was no longer required (which may have driven prefrontal activity). But in another study, PFC multi-unit activity was found to reflect an unreported feature of a stimulus, even when the animal had to report on a different, orthogonal stimulus feature (Mante et al., 2013). It is unlikely that the animal would always implicitly prepare to report on both features, given that only one report was actually required and the task was challenging with near-threshold stimuli. There was no immediate task-related benefit to represent the irrelevant feature in the brain, and yet they did so in the PFC.

**Fig. 4.**
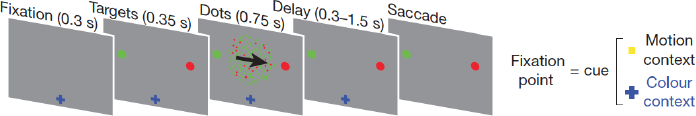
PFC activity represents specific perceptual content even when it is not relevant to current tasks. In a multi-unit recording study (Mante et al., 2013), monkeys had to make saccades to indicate either the direction or the color of the majority of moving dots, depending on the fixation cue. Task-irrelevant / unattended information (e.g., color) was decodable from neuronal activity in the frontal eye field, almost as well as task-relevant information (e.g., motion).

One could argue that because the animals switched from having to report one feature to having to report on another feature on different trials, inefficient task switching could cause the animal to implicitly prepare to report on both features immediately after a switch. Indeed, there is some evidence of the existence of such switch costs (Figure 1D & 1E in Mante et al., 2013). But such interference was relatively small in magnitude, and in general it is known that task switch effects are most prominent only during the first few trials in a block after a switch (Monsell, 2003). Here, the trials were blocked such that the animals report on the same feature for 72 consecutive trials in each block. If task switching was the main reason why the unreported feature could be decoded from PFC muti-unit activity, it would seem difficult to explain why the feature can be successfully decoded *almost to the same degree* as the attended and reported features.

This is not to say that in studies of conscious perception, the demand to make explicit reports never drives further activity in PFC. It probably does for signals obtained with most human neuroimaging methods, and this is congruent with some previous null findings, as well as the comment by Noy et al. (2015) that their positive ECoG findings in PFC were subtle when no report was required. But we emphasize that in the multi-unit recording study mentioned above (Mante et al., 2013), even unreported stimulus features could be robustly decoded to a similar degree as attended and reported features.

Overall, these findings are congruent with the typical conclusions reached by previous neuronal electrophysiology studies (Thompson and Schall, 1999, 2000) that motor report cannot be that sole reason why prefrontal neuronal activity was found to be associated with visual awareness. For example, Thompson and Schall (1999) argued that the short latencies of the relevant responses render report-based accounts unlikely. Libedinsky and Livingstone (2011) augmented their similar interpretation with careful analyses to rule out contamination by saccade generation or planning. Thompson and Schall (2000) also argued that while report-based accounts may explain the firing of movement-related neurons, they cannot account for neurons tuned specifically for visual processing. In general, these PFC neurons are thought to be involved in determining visual awareness for near-threshold visual stimuli because they contribute to the generation of the conscious percept itself.

Thus, the recent excitement surrounding the use of so-called ‘no-report’ paradigms may be misguided; PFC activity in conscious perception had already survived such tests, well before their recent advocacy.

## Concluding remarks

Consciousness is an important yet controversial topic, on which weighty claims need to be made with care. Specifically, individual null findings obtained with one measure, even if true, may not generalize to other measures. In reviewing the literature, we need to take into account research reflecting different perspectives, as well as studies conducted with different methods. When we do so, we find that the specific role of PFC in conscious perception resists even the sternest challenges.

## Acknowledgements

We would like to thank Leon Deouell, Steve Fleming, and Biyu He for their helpful comments and feedback. This was work was partially funded by NIH grant R01NS088628.

## References

Albers AM, Kok P, Toni I, Dijkerman HC, de Lange FP (2013) Shared representations for working memory and mental imagery in early visual cortex. Curr Biol 23:1427–1431.

Andersen RA, Asanuma C, Cowan WM (1985) Callosal and prefrontal associational projecting cell populations in area 7A of the macaque monkey: a study using retrogradely transported fluorescent dyes. J Comp Neurol 232:443–455.

Baars BJ (1997) In the theatre of consciousness. Global Workspace Theory, a rigorous scientific theory of consciousness. Journal of Consciousness Studies 4:292–309.

Baars BJ (2005) Global workspace theory of consciousness: toward a cognitive neuroscience of human experience. In: Progress in Brain Research (Steven Laureys, ed), pp 45–53. Elsevier.

Badre D, D’Esposito M (2009) Is the rostro-caudal axis of the frontal lobe hierarchical? Nat Rev Neurosci 10:659–669.

Barbas H, Mesulam MM (1981) Organization of afferent input to subdivisions of area 8 in the rhesus monkey. J Comp Neurol 200:407–431.

Barceló F, Knight RT (2002) Both random and perseverative errors underlie WCST deficits in prefrontal patients. Neuropsychologia 40:349–356.

Barceló F, Suwazono S, Knight RT (2000) Prefrontal modulation of visual processing in humans. Nat Neurosci 3:399–403.

Beer JS, Heerey EA, Keltner D, Scabini D, Knight RT (2003) The regulatory function of self-conscious emotion: insights from patients with orbitofrontal damage. J Pers Soc Psychol 85:594–604.

Beer JS, John OP, Scabini D, Knight RT (2006) Orbitofrontal cortex and social behavior: integrating self-monitoring and emotion-cognition interactions. J Cogn Neurosci 18:871–879.

Bhandari A, Gagne C, Badre D (2017) Just above chance: is it harder to decode information from human prefrontal cortex BOLD signals? bioRxiv:127324 Available at: http://biorxiv.org/content/early/2017/04/17/127324 [Accessed May 14, 2017].

Brascamp J, Blake R, Knapen T (2015) Negligible fronto-parietal BOLD activity accompanying unreportable switches in bistable perception. Nat Neurosci 18:1672–1678.

Brickner RM (1952) Brain of patient A. after bilateral frontal lobectomy; status of frontal-lobe problem. AMA Arch Neurol Psychiatry 68:293–313.

Buzsaki G (2006) Rhythms of the Brain. Oxford University Press.

Cavada C, Goldman-Rakic PS (1989) Posterior parietal cortex in rhesus monkey: II. Evidence for segregated corticocortical networks linking sensory and limbic areas with the frontal lobe. J Comp Neurol 287:422–445.

Chiang T-C, Lu R-B, Hsieh S, Chang Y-H, Yang Y-K (2014) Stimulation in the dorsolateral prefrontal cortex changes subjective evaluation of percepts. PLoS One 9:e106943.

Cleeremans A (2011) The Radical Plasticity Thesis: How the Brain Learns to be Conscious. Front Psychol 2:86.

Cleeremans A (2014) Connecting conscious and unconscious processing. Cogn Sci 38:1286–1315.

Cortese A, Amano K, Koizumi A, Kawato M, Lau H (2016) Multivoxel neurofeedback selectively modulates confidence without changing perceptual performance. Nat Commun 7:13669.

Croxson PL, Johansen-Berg H, Behrens TEJ, Robson MD, Pinsk MA, Gross CG, Richter W, Richter MC, Kastner S, Rushworth MFS (2005) Quantitative investigation of connections of the prefrontal cortex in the human and macaque using probabilistic diffusion tractography. J Neurosci 25:8854–8866.

de Gardelle V, Kouider S (2009) Cognitive Theories of Consciousness. Encyclopedia of consciousness 1:135–146.

Dehaene S (2014) Consciousness and the Brain: Deciphering How the Brain Codes Our Thoughts. Penguin Publishing Group.

Dehaene S, Changeux J-P (2011) Experimental and theoretical approaches to conscious processing. Neuron 70:200–227.

Dehaene S, Changeux J-P, Naccache L (2011) The Global Neuronal Workspace Model of Conscious Access: From Neuronal Architectures to Clinical Applications. In: Characterizing Consciousness: From Cognition to the Clinic? (Dehaene S, Christen Y, eds), pp 55–84 Research and Perspectives in Neurosciences. Springer Berlin Heidelberg.

Dehaene S, Changeux J-P, Naccache L, Sackur J, Sergent C (2006) Conscious, preconscious, and subliminal processing: a testable taxonomy. Trends Cogn Sci 10:204–211.

Dehaene S, Naccache L (2001) Towards a cognitive neuroscience of consciousness: basic evidence and a workspace framework. Cognition 79:1–37.

de Lafuente V, Romo R (2006) Neural correlate of subjective sensory experience gradually builds up across cortical areas. Proc Natl Acad Sci U S A 103:14266–14271.

Del Cul A, Dehaene S, Reyes P, Bravo E, Slachevsky A (2009) Causal role of prefrontal cortex in the threshold for access to consciousness. Brain 132:2531–2540.

Dienes Z (2008) Subjective measures of unconscious knowledge. Prog Brain Res 168:49–64.

Duncan J, Owen AM (2000) Common regions of the human frontal lobe recruited by diverse cognitive demands. Trends Neurosci 23:475–483.

Ester EF, Sprague TC, Serences JT (2015) Parietal and Frontal Cortex Encode Stimulus-Specific Mnemonic Representations during Visual Working Memory. Neuron 87:893–905.

Fleming SM, Ryu J, Golfinos JG, Blackmon KE (2014) Domain-specific impairment in metacognitive accuracy following anterior prefrontal lesions. Brain 137:2811–2822.

Frassle S, Sommer J, Jansen A, Naber M, Einhauser W (2014) Binocular Rivalry: Frontal Activity Relates to Introspection and Action But Not to Perception. Journal of Neuroscience 34:1738–1747.

Friedman-Hill SR, Robertson LC, Treisman A (1995) Parietal contributions to visual feature binding: evidence from a patient with bilateral lesions. Science 269:853–855.

Fulton JF (1949) Functional localization in relation to frontal lobotomy. Oxford University Press.

Giles N, Lau H, Odegaard B (2016) What Type of Awareness Does Binocular Rivalry Assess? Trends Cogn Sci 20:719–720.

Green DM, Swets JA (1966) Signal detection theory and psychophysics. 1966. New York.

Hebart MN, Schriever Y, Donner TH, Haynes J-D (2016) The Relationship between Perceptual Decision Variables and Confidence in the Human Brain. Cereb Cortex 26:118–130.

Hebb DO, Penfield W (1940) HUMAN BEHAVIOR AFTER EXTENSIVE BILATERAL REMOVAL FROM THE FRONTAL LOBES. Arch NeurPsych 44:421–438.

He BJ, Raichle ME (2009) The fMRI signal, slow cortical potential and consciousness. Trends Cogn Sci 13:302–309.

He BJ, Raichle ME (2010) The slow cortical potential hypothesis on consciousness. New Horizons in the Neuroscience of Available at: https://books.google.com/books?hl=en&lr=&id=tLMkIumdot0C&oi=fnd&pg=PA3&ots=GgqhjRSDlH&sig=ISAHN2LoX9Dyc3kxjE2IcWZh-BU.

Heekeren HR, Marrett S, Bandettini PA, Ungerleider LG (2004) A general mechanism for perceptual decision-making in the human brain. Nature 431:859–862.

Huth AG, Lee T, Nishimoto S, Bilenko NY, Vu AT, Gallant JL (2016) Decoding the Semantic Content of Natural Movies from Human Brain Activity. Front Syst Neurosci 10:81.

Jazayeri M, Movshon JA (2007) A new perceptual illusion reveals mechanisms of sensory decoding. Nature 446:912–915.

Juergens E, Guettler A, Eckhorn R (1999) Visual stimulation elicits locked and induced gamma oscillations in monkey intracortical- and EEG-potentials, but not in human EEG. Exp Brain Res 129:247–259.

Kim JN, Shadlen MN (1999) Neural correlates of a decision in the dorsolateral prefrontal cortex of the macaque. Nat Neurosci 2:176–185.

Knight RT, Grabowecky M (1995) Escape From Linear Time: Prefrontal Cortex and Conscious Experience. In: The Cognitive Neurosciences (Gazzaniga MS, ed). MIT Press.

Koch C, Massimini M, Boly M, Tononi G (2016a) Neural correlates of consciousness: progress and problems. Nat Rev Neurosci 17:307–321.

Koch C, Massimini M, Boly M, Tononi G (2016b) Posterior and anterior cortex [mdash] where is the difference that makes the difference? Nat Rev Neurosci Available at: http://dx.doi.org/10.1038/nrn.2016.105 [Accessed July 30, 2016].

Komura Y, Nikkuni A, Hirashima N, Uetake T, Miyamoto A (2013) Responses of pulvinar neurons reflect a subject’s confidence in visual categorization. Nat Neurosci 16:749–755.

Kouider S, de Gardelle V, Sackur J, Dupoux E (2010) How rich is consciousness? The partial awareness hypothesis. Trends Cogn Sci 14:301–307.

Kouider S, Dehaene S, Jobert A, Le Bihan D (2007) Cerebral bases of subliminal and supraliminal priming during reading. Cereb Cortex 17:2019–2029.

Ko Y, Lau H (2012) A detection theoretic explanation of blindsight suggests a link between conscious perception and metacognition. Philos Trans R Soc Lond B Biol Sci, 367:1401–1411.

Lak A, Costa GM, Romberg E, Koulakov AA, Mainen ZF, Kepecs A (2014) Orbitofrontal cortex is required for optimal waiting based on decision confidence. Neuron 84:190–201.

Lamme VAF (2003) Why visual attention and awareness are different. Trends Cogn Sci 7:12–18.

Lamme VAF (2006) Towards a true neural stance on consciousness. Trends Cogn Sci 10:494–501.

Lamme VAF (2010) How neuroscience will change our view on consciousness. Cogn Neurosci 1:204–220.

Lau HC (2008) Are we studying consciousness yet? In: Frontiers of Consciousness. Oxford: Oxford University Press.

Lau H, Rosenthal D (2011) Empirical support for higher-order theories of conscious awareness. Trends Cogn Sci 15:365–373.

Lee TG, D’Esposito M (2012) The dynamic nature of top-down signals originating from prefrontal cortex: a combined fMRI-TMS study. J Neurosci 32:15458–15466.

Libedinsky C, Livingstone M (2011) Role of prefrontal cortex in conscious visual perception. J Neurosci 31:64–69.

Lumer ED, Rees G (1999) Covariation of activity in visual and prefrontal cortex associated with subjective visual perception. Proc Natl Acad Sci U S A 96:1669–1673.

Mackey WE, Devinsky O, Doyle WK, Meager MR, Curtis CE (2016) Human Dorsolateral Prefrontal Cortex Is Not Necessary for Spatial Working Memory. J Neurosci 36:2847–2856.

Maniscalco B, Lau H (2012) A signal detection theoretic approach for estimating metacognitive sensitivity from confidence ratings. Conscious Cogn 21:422–430.

Mante V, Sussillo D, Shenoy KV, Newsome WT (2013) Context-dependent computation by recurrent dynamics in prefrontal cortex. Nature 503:78–84.

McCurdy LY, Maniscalco B, Metcalfe J, Liu KY, de Lange FP, Lau H (2013) Anatomical coupling between distinct metacognitive systems for memory and visual perception. J Neurosci 33:1897–1906.

Melnick MD, Tadin D, Huxlin KR (2016) Relearning to See in Cortical Blindness. Neuroscientist 22:199–212.

Merikle PM, Smilek D, Eastwood JD (2001) Perception without awareness: perspectives from cognitive psychology. Cognition 79:115–134.

Mettler FA (1949) Selective partial ablation of the frontal cortex, a correlative study of its effects on human psychotic subjects. Hoeber.

Miller EK (2000) The prefrontal cortex and cognitive control. Nat Rev Neurosci 1:59–65.

Miller EK, Cohen JD (2001) An integrative theory of prefrontal cortex function. Annu Rev Neurosci 24:167–202.

Miyamoto K, Osada T, Setsuie R, Takeda M, Tamura K, Adachi Y, Miyashita Y (2017) Causal neural network of metamemory for retrospection in primates. Science 355:188–193.

Monsell S (2003) Task switching. Trends Cogn Sci 7:134–140.

Nakamura RK, Mishkin M (1986) Chronic “blindness” following lesions of nonvisual cortex in the monkey. Exp Brain Res 63:173–184.

Niedermeyer E, da Silva FHL (2005) Electroencephalography: Basic Principles, Clinical Applications, and Related Fields. Lippincott Williams & Wilkins.

Niendam TA, Laird AR, Ray KL, Dean YM, Glahn DC, Carter CS (2012) Meta-analytic evidence for a superordinate cognitive control network subserving diverse executive functions. Cogn Affect Behav Neurosci 12:241–268.

Noy N, Bickel S, Zion-Golumbic E, Harel M, Golan T, Davidesco I, Schevon CA, McKhann GM, Goodman RR, Schroeder CE, Mehta AD, Malach R (2015) Ignition’s glow: Ultra-fast spread of global cortical activity accompanying local “ignitions” in visual cortex during conscious visual perception. Conscious Cogn 35:206–224.

Panagiotaropoulos TI, Deco G, Kapoor V, Logothetis NK (2012) Neuronal discharges and gamma oscillations explicitly reflect visual consciousness in the lateral prefrontal cortex. Neuron 74:924–935.

Passingham RE, Wise SP (2012) The Neurobiology of the Prefrontal Cortex: Anatomy, Evolution, and the Origin of Insight. OUP Oxford.

Perry A, Lwi SJ, Verstaen A, Dewar C, Levenson RW, Knight RT (2016) The role of the orbitofrontal cortex in regulation of interpersonal space: evidence from frontal lesion and frontotemporal dementia patients. Soc Cogn Affect Neurosci 11:1894–1901.

Persaud N, Davidson M, Maniscalco B, Mobbs D, Passingham RE, Cowey A, Lau H (2011) Awareness-related activity in prefrontal and parietal cortices in blindsight reflects more than superior visual performance. Neuroimage 58:605–611.

Pessoa L, Adolphs R (2010) Emotion processing and the amygdala: from a “low road” to “many roads” of evaluating biological significance. Nat Rev Neurosci 11:773–783.

Petrides M, Pandya DN (1984) Projections to the frontal cortex from the posterior parietal region in the rhesus monkey. J Comp Neurol 228:105–116.

Philiastides MG, Auksztulewicz R, Heekeren HR, Blankenburg F (2011) Causal role of dorsolateral prefrontal cortex in human perceptual decision making. Curr Biol 21:980–983.

Pitts MA, Padwal J, Fennelly D, Martínez A, Hillyard SA (2014) Gamma band activity and the P3 reflect post-perceptual processes, not visual awareness. Neuroimage 101:337–350.

Pollen DA (1999) On the neural correlates of visual perception. Cereb Cortex 9:4–19.

Rahnev D, Nee DE, Riddle J, Larson AS, D’Esposito M (2016) Causal evidence for frontal cortex organization for perceptual decision making. Proc Natl Acad Sci U S A 113:6059–6064.

Ratcliff R (1978) A theory of memory retrieval. Psychol Rev 85:59.

Richter FR, Cooper RA, Bays PM, Simons JS (2016) Distinct neural mechanisms underlie the success, precision, and vividness of episodic memory. Elife 5 Available at: http://dx.doi.org/10.7554/eLife.18260.

Rigotti M, Barak O, Warden MR, Wang X-J, Daw ND, Miller EK, Fusi S (2013) The importance of mixed selectivity in complex cognitive tasks. Nature 497:585–590.

Ritzinger B, Huberle E, Karnath H-O (2012) Bilateral theta-burst TMS to influence global gestalt perception. PLoS One 7:e47820.

Romanski LM, Giguere M, Bates JF, Goldman-Rakic PS (1997) Topographic organization of medial pulvinar connections with the prefrontal cortex in the rhesus monkey. J Comp Neurol 379:313–332.

Rounis E, Maniscalco B, Rothwell JC, Passingham RE, Lau H (2010) Theta-burst transcranial magnetic stimulation to the prefrontal cortex impairs metacognitive visual awareness. Cogn Neurosci 1:165–175.

Ruff CC, Blankenburg F, Bjoertomt O, Bestmann S, Freeman E, Haynes J-D, Rees G, Josephs O, Deichmann R, Driver J (2006) Concurrent TMS-fMRI and psychophysics reveal frontal influences on human retinotopic visual cortex. Curr Biol 16:1479–1488.

Sandberg K, Frässle S, Pitts M (2016) Future directions for identifying the neural correlates of consciousness. Nat Rev Neurosci Available at: http://dx.doi.org/10.1038/nrn.2016.104.

Shipp S (2003) The functional logic of cortico-pulvinar connections. Philos Trans R Soc Lond B Biol Sci 358:1605–1624.

Siclari F, Baird B, Perogamvros L, Bernardi G, LaRocque JJ, Riedner B, Boly M, Postle BR, Tononi G (2017) The neural correlates of dreaming. Nat Neurosci Available at: http://dx.doi.org/10.1038/nn.4545.

Stokes MG (2015) “Activity-silent” working memory in prefrontal cortex: a dynamic coding framework. Trends Cogn Sci 19:394–405.

Stuss DT, Knight RT (2013) Principles of Frontal Lobe Function. OUP USA.

Szczepanski SM, Knight RT (2014) Insights into human behavior from lesions to the prefrontal cortex. Neuron 83:1002–1018.

Thompson KG, Schall JD (1999) The detection of visual signals by macaque frontal eye field during masking. Nat Neurosci 2:283–288.

Thompson KG, Schall JD (2000) Antecedents and correlates of visual detection and awareness in macaque prefrontal cortex. Vision Res 40:1523–1538.

Tononi G, Boly M, Massimini M, Koch C (2016) Integrated information theory: from consciousness to its physical substrate. Nat Rev Neurosci 17:450–461.

Tse PU, Martinez-Conde S, Schlegel AA, Macknik SL (2005) Visibility, visual awareness, and visual masking of simple unattended targets are confined to areas in the occipital cortex beyond human V1/V2. Proc Natl Acad Sci U S A 102:17178–17183.

Tsuchiya N, Wilke M, Frässle S, Lamme VAF (2015) No-Report Paradigms: Extracting the True Neural Correlates of Consciousness. Trends Cogn Sci 19:757–770.

Turatto M, Sandrini M, Miniussi C (2004) The role of the right dorsolateral prefrontal cortex in visual change awareness. Neuroreport 15:2549–2552.

Voytek B, Davis M, Yago E, Barceló F, Vogel EK, Knight RT (2010) Dynamic neuroplasticity after human prefrontal cortex damage. Neuron 68:401–408.

Wang M, Arteaga D, He BJ (2013) Brain mechanisms for simple perception and bistable perception. Proc Natl Acad Sci U S A 110:E3350–E3359.

Weiskrantz L (1997) Consciousness Lost and Found: A Neuropsychological Exploration. Oxford University Press.

